## Supplementary figures and images for "The Mac1 ADP-ribosylhydrolase is a Therapeutic Target for SARS-CoV-2"

### Actin.tif

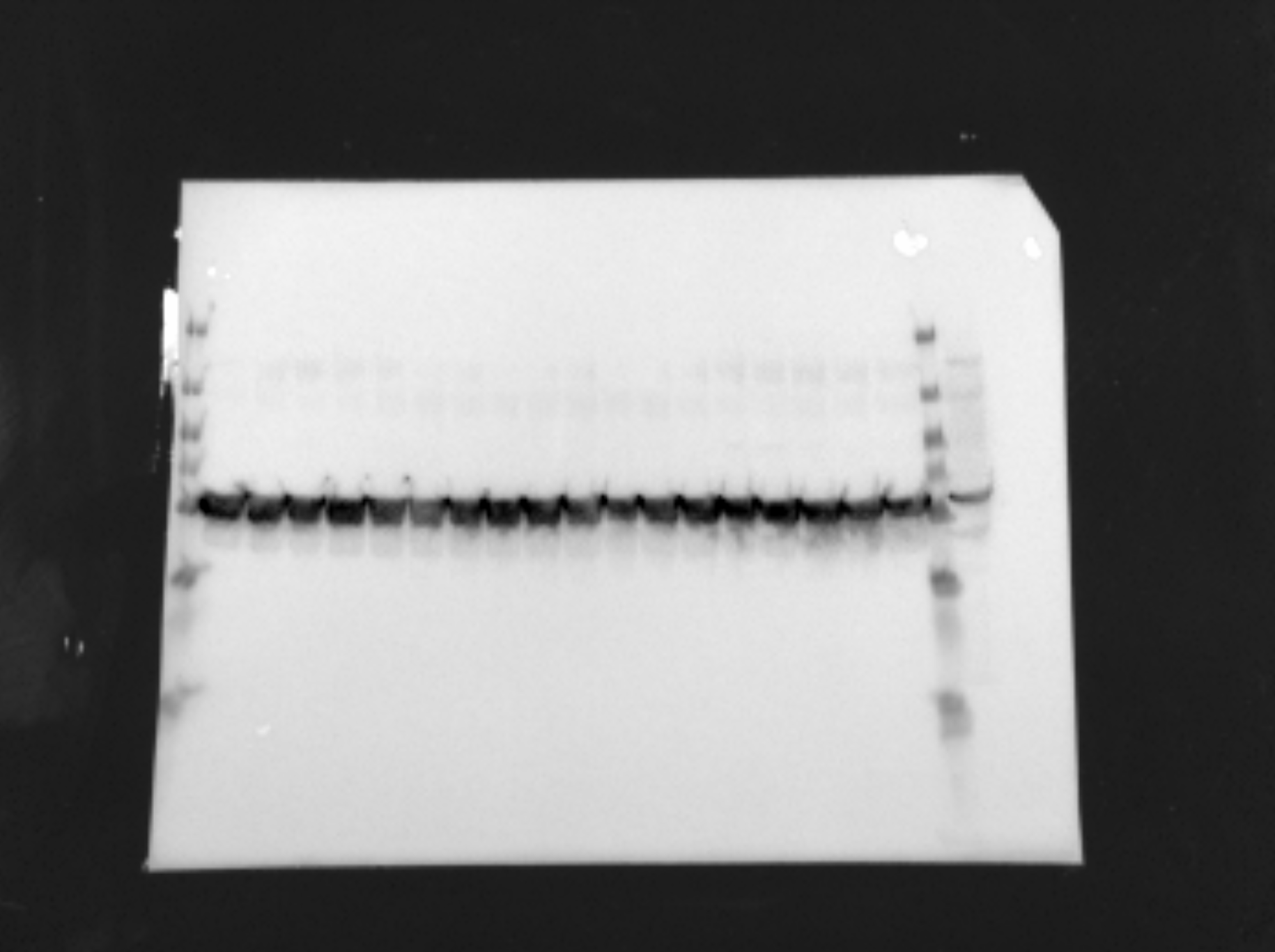

### Ashworth 2024-04-09 11h16m17s(Chemiluminescence).raw16.tif

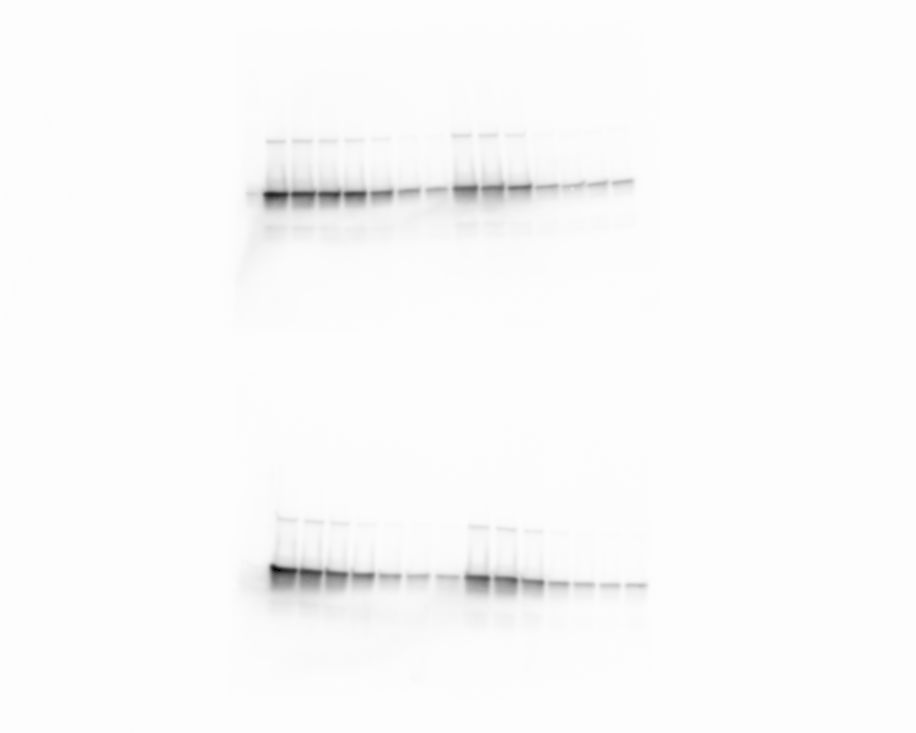

### Ashworth 2024-04-09 11h21m29s(Colorimetric).tif

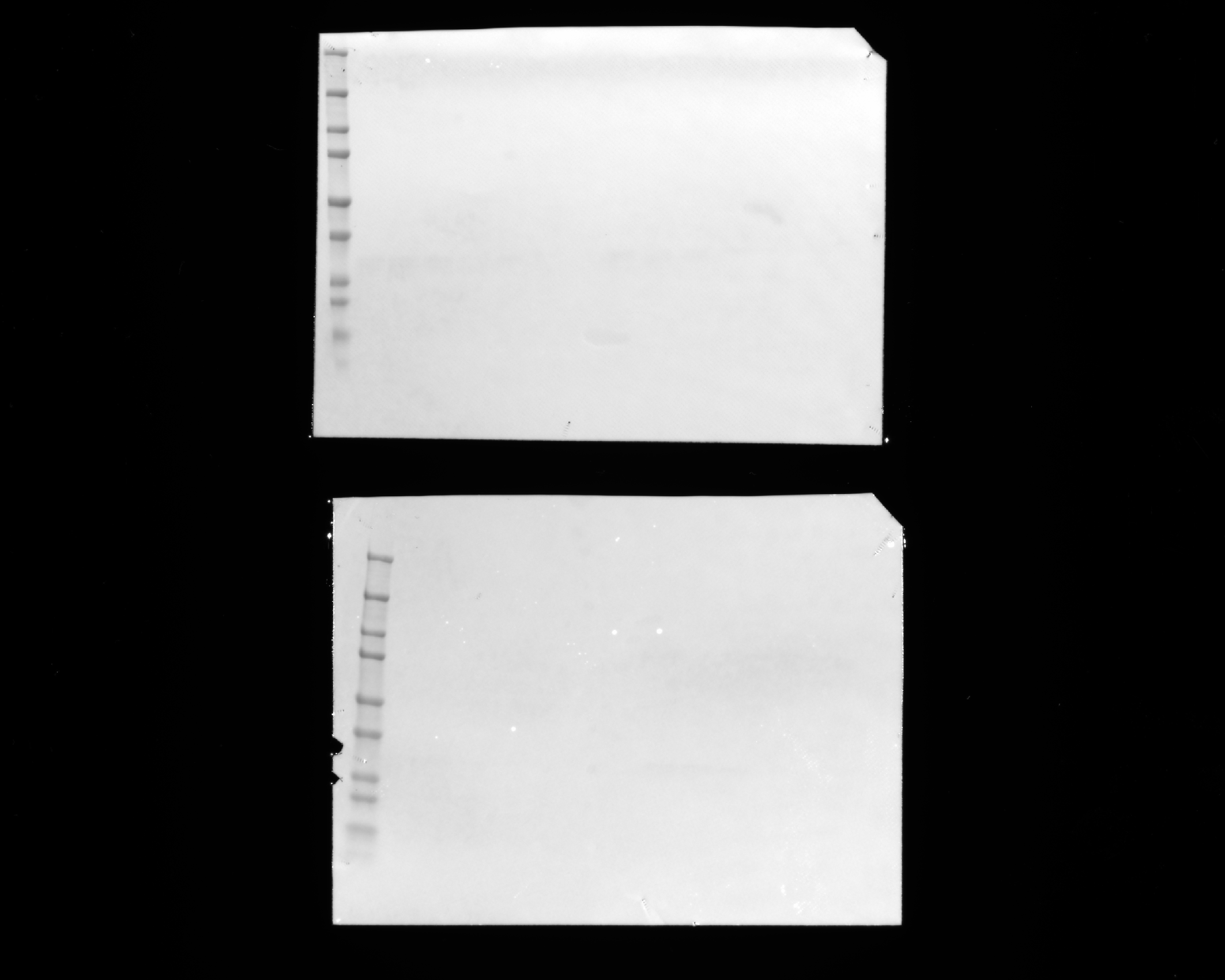

### Figure 2 - Figure Supplement 1 - source data 1

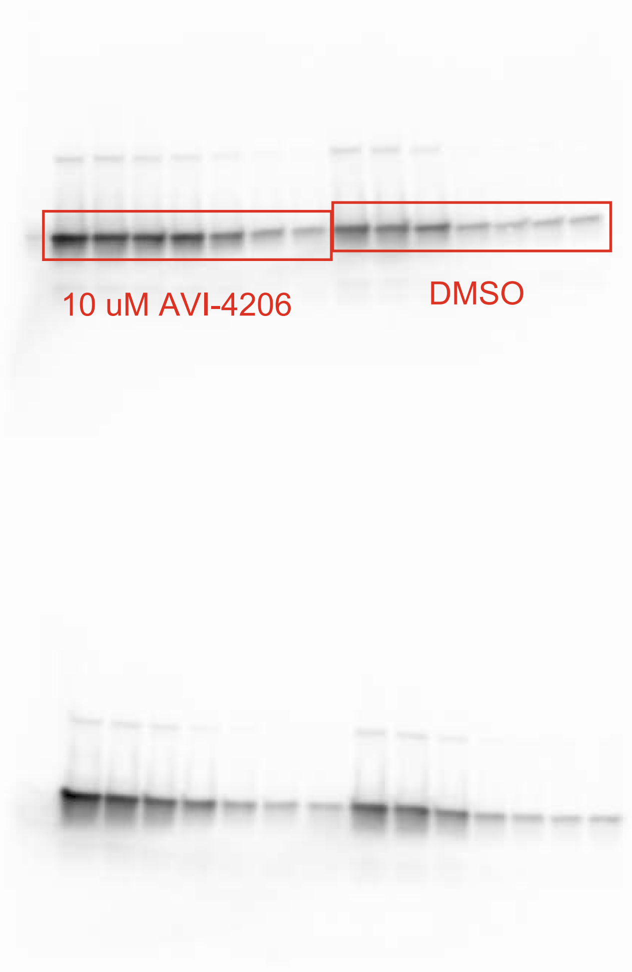

### Figure 3 - Figure Supplement 3 - source data 1

A

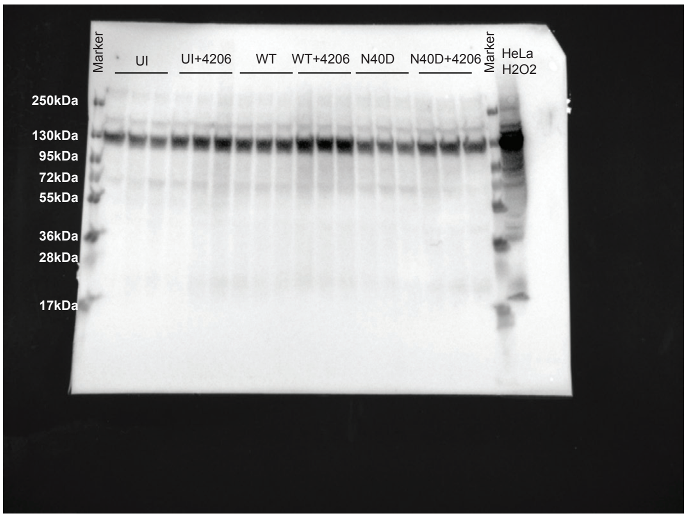


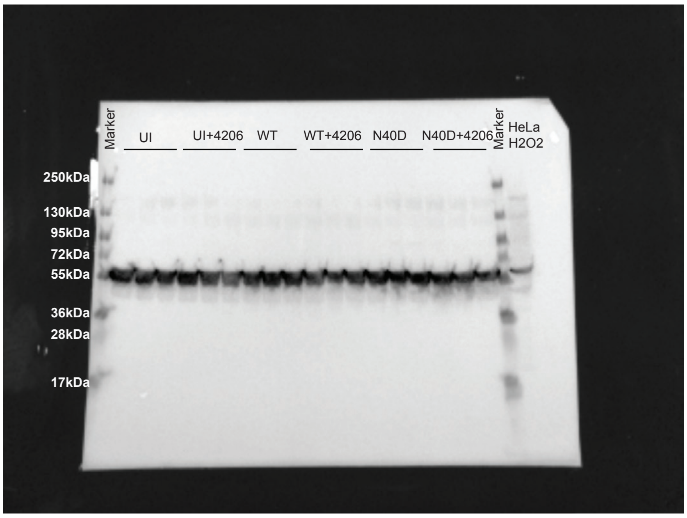


C

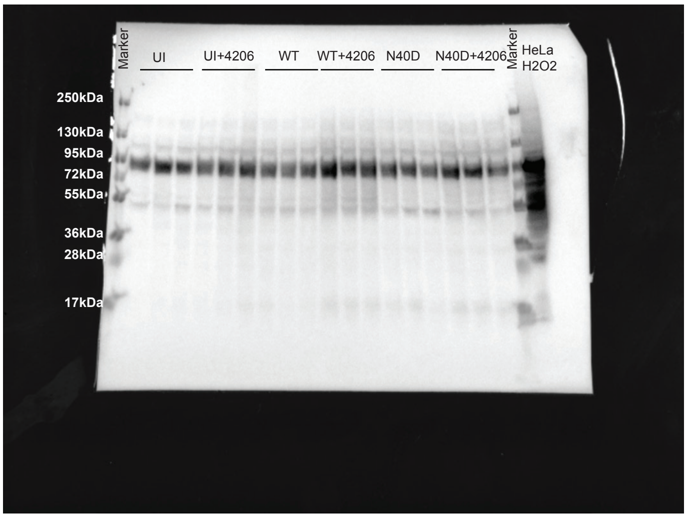


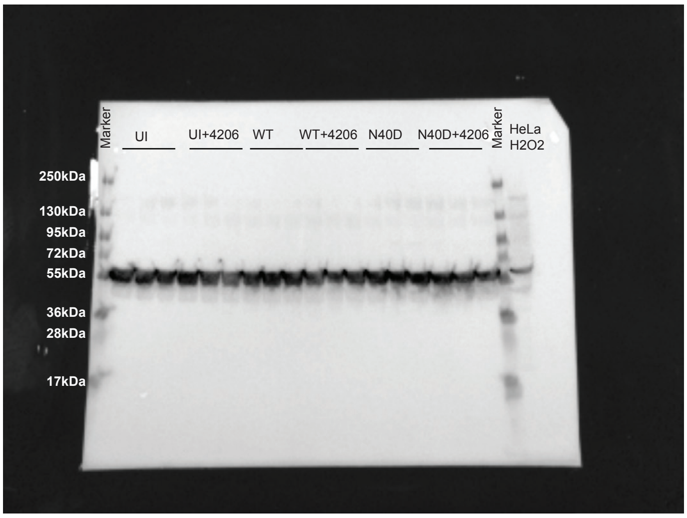


**Figure 3-figure supplement 3-source data 2**

### MonoADpr.tif

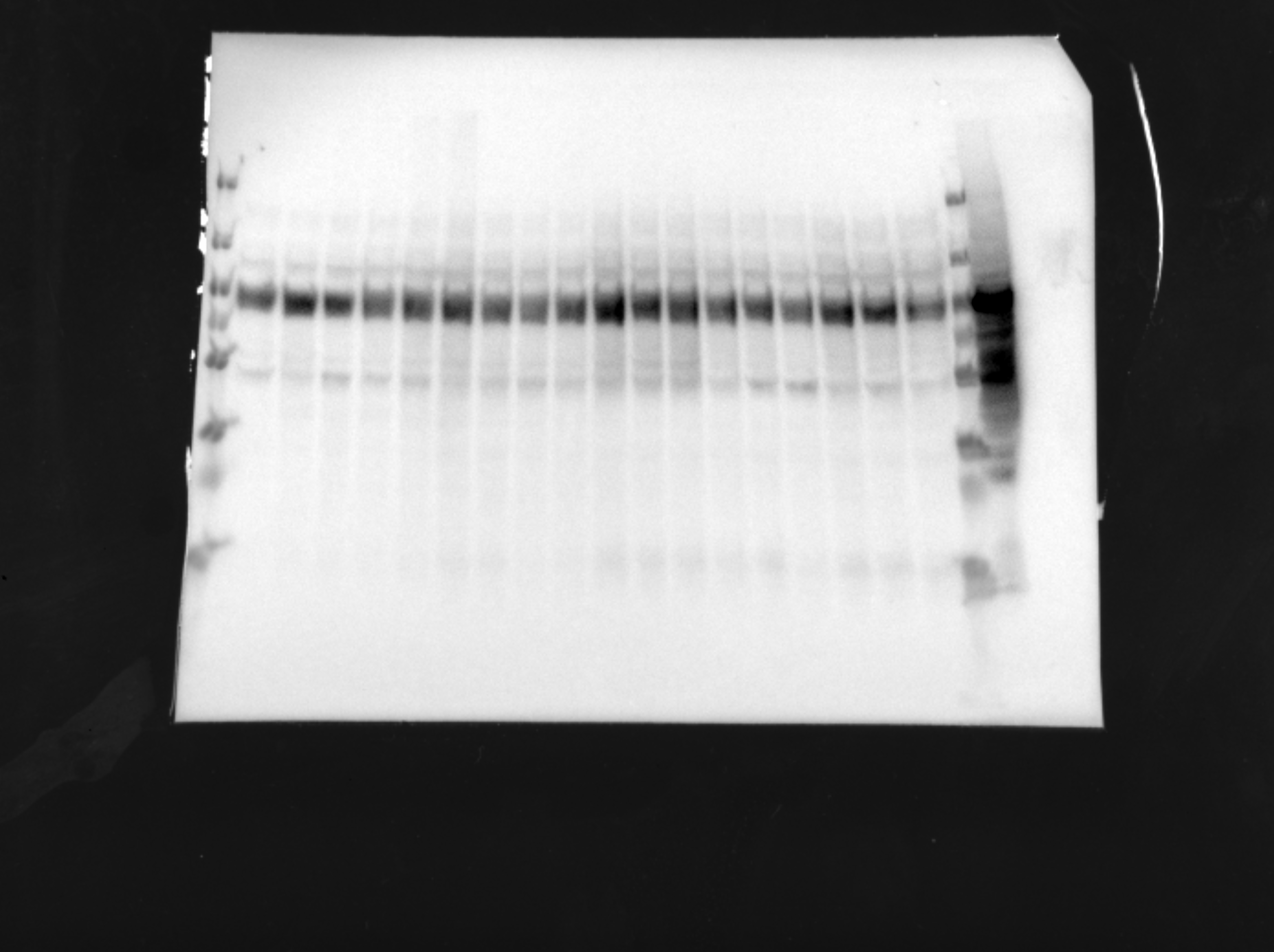

### pan-ADPr.png

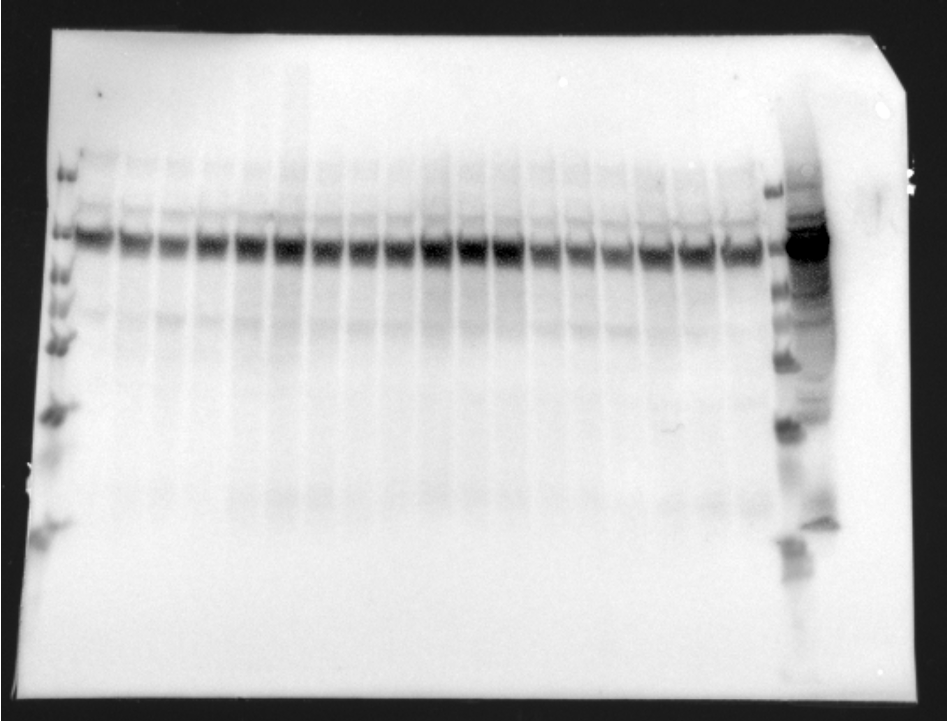
